# How forgiving are M/EEG inverse solutions to noise level misspecification? An excursion into the BSI-Zoo

**DOI:** 10.1101/2025.03.12.642831

**Authors:** Anuja Negi, Stefan Haufe, Alexandre Gramfort, Ali Hashemi

## Abstract

Brain source imaging (BSI), also known as source localization, from magneto- and electroencephalographic (M/EEG) data, is a challenging ill-posed inverse problem. Accurate source estimation is sensitive to modeling parameters, such as regularization strength and noise level, where misconfigurations can lead to under- or overfitting. Different BSI methods, however, may vary in their robustness to suboptimal parameter choices. Here we conducted extensive simulations of brain sources superimposed by varying degrees of sensor noise to study the ranges of noise misspecification within which different BSI approaches can still localize well. Using the Earth Mover’s Distance (EMD) and other metrics, we compare the performance of smooth linear inverse solutions with that of sparse non-linear Bayesian learning solutions. Additionally, we assess the effectiveness of various noise estimation and cross-validation techniques to select hyperparameters close to those achieving optimal localization. Methods and experiments are made available within the BSI-Zoo Python package.

## 1 Introduction

Electroencephalography (EEG) and magnetoencephalography (MEG) are non-invasive techniques used to capture the brain’s electrical activity with high temporal resolution, serving as essential tools in both fundamental investigation in neuroscience and application within clinical neurology. Brain Source Imaging (BSI) using M/EEG data, also known as the E/MEG inverse problem or brain source localization, involves reconstructing and localizing neural activation from recorded M/EEG signals. This task is inherently challenging because it represents an ill-posed inverse problem: the number of sensors is typically far fewer than the possible brain source locations, resulting in multiple potential solutions that can fit the observed data equally well. Additionally, the mapping between brain sources and sensor recordings is often highly unstable, which complicates accurate source localization and limits spatial resolution [1, 2, 3]. To address these issues, a variety of source reconstruction techniques have been developed, incorporating prior assumptions to improve localization accuracy and reliability.

Classical approaches such as the minimum-norm estimates use Gaussian priors to promote smoothness in the solution [1, 2, 3], while Lasso-based methods employ 𝓁_1_-norm penalties to encourage sparsity, making them suitable for identifying focal brain activity related to specific cognitive tasks [4]. Extending this, techniques such as the Mixed-Norm Estimate (MxNE) [5], introduce structured sparsity across spatial and temporal domains by utilizing mixed norms. Techniques like S-FLEX [6, 7, 8], ir-MxNE [9], TF-MxNE [10], irTF-MxNE [11], and STOUT [12] enforce sparsity in specific domains, such as cortical patches or Gabor frames, through appropriate norm constraints.

These approaches are typically classified as Maximum-A-Posteriori (MAP) methods, as they assume fixed prior distributions. However, there are situations where it is beneficial to explore entire families of distributions defined by a set of hyperparameters. These hyperparameters can be learned from the data alongside the model parameters using a hierarchical empirical Bayesian approach, where the hyperparameters are estimated through the Maximum Likelihood (ML) principle. Various instantiations of this approach have been introduced under different names, such as Sparse Bayesian Learning (SBL) [13], Automatic Relevance Determination (ARD) [14], Kernel Fisher Discriminant (KFD) [15], and Variational Bayes (VB) [16, 17]. For a comprehensive overview of Bayesian machine learning techniques for EEG/MEG signals, readers can refer to [18].

While these various methods originate from different backgrounds, they share a fundamental requirement for parameter tuning. The accuracy of brain source localization heavily relies on precise adjustment of modeling parameters, such as regularization strength and noise level estimation, to balance model fit and complexity. These hyperparameters play a critical role in avoiding both overfitting and underfitting and thus, their selection strongly impacts the accuracy of BSI results (as illustrated in 4a).

Furthermore, it is noteworthy that different BSI methods exhibit inherent variations in their ability to localize in different signal-to-noise ratio regimes and for different amounts of regularization. Understanding how various BSI techniques are differentially affected by noise level parameter specification in terms of their resulting bias-variance tradeoff is essential for attaining accurate source localization performance.

Despite constant advancement of BSI methodologies, tools and frameworks that support consistent parameter tuning and reproducibility across methods are scarce. Specifically, there is a lack of open-source Python packages offering a comprehensive suite of state-of-the-art BSI methods, limiting accessibility for researchers seeking a unified platform for method comparison. Existing packages lack the diversity of methods or the flexibility needed for a robust analysis of BSI techniques, and they rarely provide resources for reproducing code, making it difficult to validate results or build upon prior work [19]. Additionally, the absence of an easy-to-use experimental framework prevents researchers from efficiently testing and benchmarking BSI methods under consistent conditions. Current software options are typically complex, requiring significant customization, and are primarily designed to support neuroscientific studies using published methods rather than facilitating comparative studies, particularly those involving large-scale simulations. As a result, they lack built-in functionality for systematically benchmarking different approaches under controlled conditions. Finally, existing studies do not quantify the impact of noise level misspecification on localization performance, often relying on heuristic approaches to fix regularization parameters inconsistently across methods and reporting results based on fixed regularization. This overlooks the potential for noise misspecification, a critical factor that can lead to overfitting or underfitting in practice. Addressing these gaps is crucial for developing robust BSI applications and advancing the field.

Our contributions to this paper address these challenges and are threefold:

- As our first contribution, we develop a Python package, named “BSI-Zoo”^1^, serving as a unifying platform for the comparative analysis of various BSI approaches. This open-source package is intended to facilitate the assessment and comparison of innovations in brain source imaging research.
- Second, we design and implement a comprehensive simulation environment that introduces different levels of sensor noise. This environment builds on the BSI-Zoo package and enables the controlled testing of BSI methods under varying conditions. By simulating various degrees of sensor noise, we can investigate the robustness of different approaches within the BSI domain.
- Third, using the developed simulation environment, we conduct a rigorous evaluation of distinct approaches from two fundamental paradigms within the BSI domain: smooth linear inverse solutions and sparse non-linear Bayesian learning solutions. Our study focuses on learning about noise and applying cross-validation techniques to identify the elusive optimal points for source localization accuracy. This investigation defines the limits within which different BSI approaches can sustain their capability for accurate source localization and provides valuable insights to improve their performance.

## 2 Methodology

### 2.1 Brain Source Imaging Forward Model

The E/MEG measurement process is described by the linear *forward model*: **Y** = **LX** + **E**. This model maps the electrical activity originating from brain sources, denoted as **X**, to the measurements obtained from sensors, denoted as **Y**. The *measurement matrix* **Y** ∈ ℝ^*M ×T*^ captures the activity recorded by *M* sensors over *T* time instances, where **y**(*t*) ∈ ℝ^*M*^, *t* = 1, …, *T*. In the case of EEG, these sensors are placed on the scalp, whereas for MEG, the sensors are positioned around the head to detect the magnetic fields generated by neural activity. The *source matrix* **X** ∈ ℝ^*N ×T*^ represents the unknown activity of *N* brain sources situated in the cortical gray matter during the same time instances, with **x**(*t*) ∈ ℝ^*N*^, *t* = 1, …, *T*. The matrix **E** = [**e**(1), …, **e**(*T*)] ∈ ℝ^*M ×T*^ accounts for *T* time points of Independent and Identically Distributed (I.I.D) zero-mean white Gaussian noise, each with a variance of *σ*^2^. This noise is represented as **e**(*t*) ∈ ℝ^*M*^ ∼ 𝒩 (0, *σ*^2^**I**_*M*_), *t* = 1, …, *T*, and is assumed to be independent of the source activities. Based on these assumptions, the likelihood of the measurements **Y** given the source activity **X** is given by:

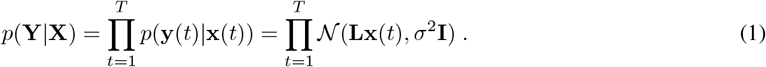

Note that the assumption of uncorrelated noise between sensors can be considered unrealistic, although this is typically handled by a whitening step discussed below [20].

### 2.2 Type-I Bayesian Learning

To address the ill-posed nature of the E/MEG inverse problem, one of the primary approaches is Maximum-A-Posteriori (MAP) estimation, commonly referred to as Bayesian Type-I learning. In this approach, a prior distribution, *p*(**X**), is assumed for the source activity to model various characteristics of brain sources, such as spatial and temporal structures. By applying Bayes’ formula, the combination of this prior with the data likelihood yields the posterior distribution of the sources given the observed data **Y**, denoted *p*(**X** | **Y**). The MAP estimate identifies the source configuration that maximizes this posterior distribution, balancing the observed data with prior assumptions about the sources. Formally, the MAP estimate is defined as:

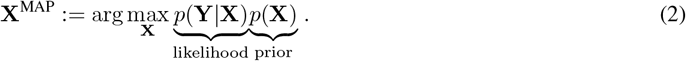

This formulation leads to an optimization problem, where the MAP estimate can equivalently be expressed as:

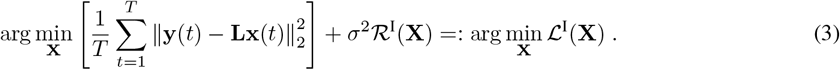

Here, ℛ^I^(**X**) = −log (*p*(**X**)) and ℒ^I^(**X**) represents the objective functions of Type-I Bayesian learning.

Through the Type-I Bayesian learning framework, the prior, *p*(**X**), provides essential guidance to align the solution with physiologically plausible characteristics of brain sources, while the observed data ensures accurate localization. Various algorithms in the BSI domain are distinguished by their choice of prior distribution, which models different assumptions about brain sources. For instance, selecting a Gaussian prior leads to the classical minimum-norm estimate [1, 2, 3], also known as the 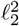-norm (or Tikhonov) regularization and referred to as “ridge regression” in statistics and machine learning. Alternatively, using a Laplace prior yields the minimum-current estimate [4], commonly known as 𝓁_1_-norm regularization or “LASSO” regression. This paper specifically focuses on implementing three different variants of sparse iterative methods and eLORETA [2], a smooth linear solver, all of which fall under the category of Type-I Bayesian methods. These methods will be described in detail in the following subsections.

#### 2.2.1 eLORETA

Exact Low-Resolution Electromagnetic Tomography (eLORETA), introduced by Pascual-Marqui [3], is a source localization method designed to provide unbiased inverse solutions by addressing depth bias inherent in M/EEG inverse problems. eLORETA addresses the issue of depth bias, which often affects localization accuracy, by introducing a depth-weighted regularization scheme that ensures consistent localization performance across brain regions at varying depths. eLORETA achieves this by minimizing the following objective function:

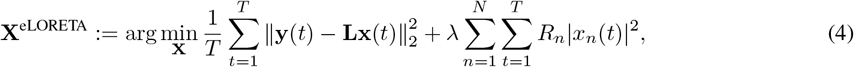

where *λ >* 0 is the regularization parameter controlling the balance between fitting the observed data and enforcing the prior, and *x*_*n*_(*t*) represents the source activity for source *n* at time *t*. The term 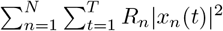 acts as a regularization prior that penalizes the norm of the sources, effectively discouraging unreasonably large amplitudes for deeper sources that would otherwise dominate the solution.

Unlike traditional fixed-parameter regularization methods, eLORETA iteratively updates the regularization weights *R*_*n*_ to adaptively compensate for depth bias in the lead field matrix, which underrepresents the contributions of deeper sources. This adaptive reweighting ensures a balanced, physiologically meaningful estimation, yielding source estimates that are both depth-unbiased and neurophysiologically plausible. To account for sensor noise and improve stability, the lead field matrix **L** is pre-whitened, resulting in the matrix **G** = **WL**, where **W** is the whitening matrix derived from the noise covariance matrix. This pre-whitening ensures isotropic noise handling and stabilizes the solution [20].

The eLORETA algorithm follows an iterative procedure, where at each iteration *k*, the regularization matrix **R** is updated to refine the inverse solution. Before solving for the sources **X** in Eq. 4, we first describe the iterative update of **R**. This matrix is initialized as an identity matrix (i.e. **R**_0_ = **I**), ensuring an unbiased starting point.

At each iteration, the regularization weights *R*_*n*_ are updated using the following rule

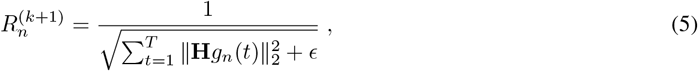

where **H** is derived from the singular value decomposition (SVD) of **GRG**^⊤^, and **g**_*n*_(*t*) represents the contribution of source *n* at time *t*. The parameter *ϵ* ensures numerical stability during the iterative process.

Convergence is determined by monitoring the relative change in **R** between iterations, defined as 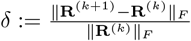. The iterations stop when δ *< τ*, where the threshold *τ* is a small predefined value.

Once the iterative process has converged, the eLORETA kernel is computed as:

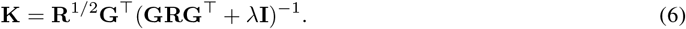

The final source estimate is derived by applying the kernel to the pre-whitened sensor measurements: **X** = **KY**.

#### 2.2.2 Sparse Iterative Type-I Methods

The sparse Type-I iterative methods, introduced in the field of BSI by [5] and [9], promote structured sparsity by incorporating a non-convex regularization term, 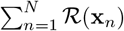, to capture the characteristics of brain source activities. The general objective function for these methods can be expressed as:

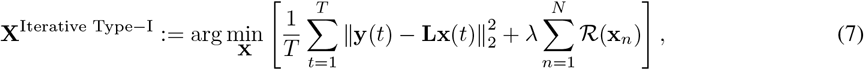

where ℛ(**x**_*n*_) is the regularization function tailored to promote sparsity, and *λ* is a regularization parameter that controls the trade-off between data fidelity and sparsity. This objective function is minimized iteratively by solving a sequence of weighted norm problems as follows:

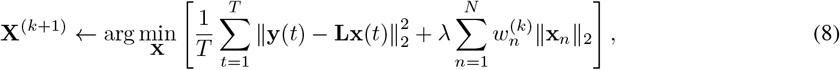

The weights 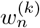 are iteratively updated based on the temporal energy of each source **x**_*n*_, defined as 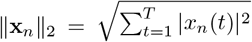. The regularization function ℛ(**x**_*n*_) and weight update 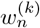 vary according to the chosen iterative method. Below, we present the specific regularization functions ℛ(**x**_*n*_) and iterative weight updates 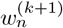 for the methods implemented in this paper.

- **Iterative** 𝓁_1_ [21]:

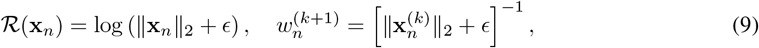

where 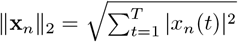
- **Iterative** 𝓁_2_ [21]:

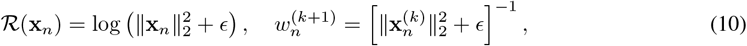

where 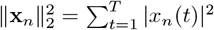
- **Iterative** 𝓁_0.5_ [9]:

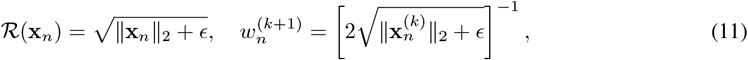

which is equivalent to

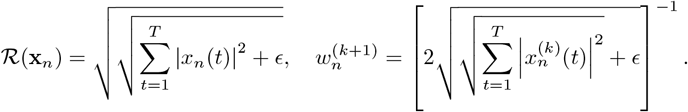

In each case, *ϵ* is a small constant added to ensure numerical stability.

### 2.3 Type-II Bayesian Learning

While the MAP technique maintains a fixed prior distribution, it is often beneficial to consider entire families of distributions denoted as *p*(**X**|***γ***), which are parameterized by a set of hyperparameters *γ*. These hyperparameters can be learned from the data concurrently with the model parameters using the Type-II Maximum Likelihood (ML-II, or simply Type-II) approach [15, 13, 22]. In the Type-II approach, the hyper-parameters are estimated using the Maximum Likelihood principle [22]:

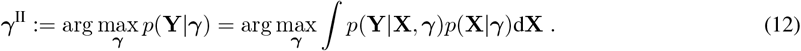

Formally, the conditional density *p*(**Y**|***γ***) is determined by integrating across all possible source distributions for a specific selection of ***γ***. The solution that maximizes Eq.(12) subsequently defines a prior distribution that is data-driven, denoted as *p*(**X**|***γ***^II^). When incorporated into the MAP estimation framework as depicted in Eq.(2), this process leads to the emergence of the Type-II source estimate, referred to as **X**^II^.

#### 2.3.1 Champagne or *γ*−MAP

Sparse Bayesian Learning (SBL), also known as the ***γ***-MAP, is a concrete instantiation of the Type-II Bayesian learning approach. In the field of brain source imaging, it has been further introduced under the name Champagne [16]. Champagne models the current activity of the brain sources as Gaussian scale mixtures: 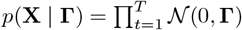, where **Γ** denotes the source covariance with a diagonal structure **Γ** = diag(***γ***) = diag([*γ*_1_, …, *γ*_*N*_]^⊤^).

Given this assumption, the posterior source distribution:

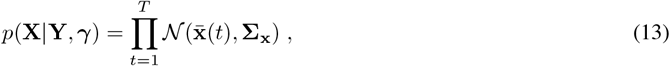

where

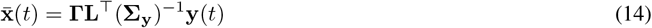

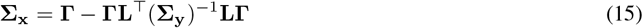

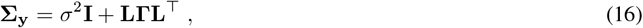

is derived as a result of learning **Γ** (hyper-parameters) through minimizing the negative log-likelihood (Type-II) loss, − log *p*(**Y** | **Γ**):

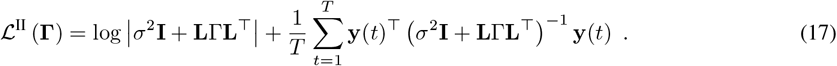

#### 2.3.2 Sparse Iterative Type-II Methods

As shown in [23, 24], the SBL cost function in Eq. 17 can be equivalently reformulated as ℒ^II−*x*^(**X, *γ***) by utilizing mathematical tools grounded in Legendre-Fenchel duality theory [25, 26]. The optimizers, {***γ***^*^, **X**^*^} are then obtained through a joint minimization over **X** and ***γ*** as

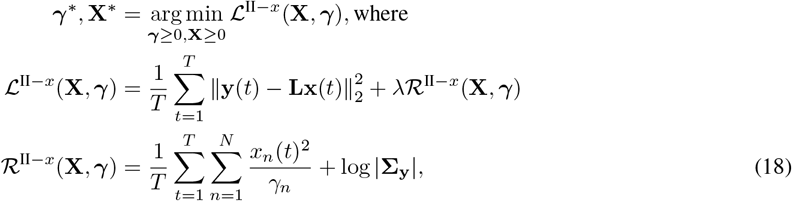

where ℛ^II−*x*^(**X, *γ***) serves as a regularizer dependent on the data **x**(*t*), with *x*_*n*_(*t*) indicating the activity of source *n* at time instant *t*. This formulation gives another view on Type-II methods in the original inverse parameter space, also noted as **x**-based space, rather than the hyperparameter space, or *γ*-based space, represented in Eq 17.

By utilizing Eq. 18, we can integrate Type-II variants of sparse iterative methods, originally proposed in [21]. In these variants, the Type-II counterparts of the iterative-𝓁_1_ and iterative-𝓁_2_ methods are derived by selecting 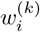 as follows:

- **Type-II Iterative** 𝓁_1_ [21]:

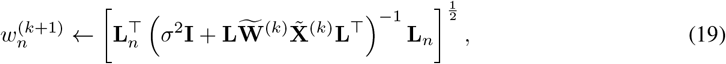
- **Type-II Iterative** 𝓁_2_ [21]:

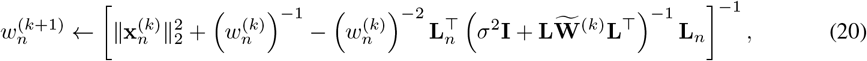

where 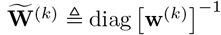 is the inverse diagonal matrix formed from the weight vector **w**^(*k*)^, which consists of elements 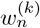 for *n* = 1, …, *N*. Similarly, 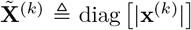 is the diagonal matrix formed from the absolute values of the source estimates at the current iteration. Finally, **L**_*n*_ denotes the *n*-th column of the lead field matrix **L**, representing the contribution of source *n* to the measurements.

##### Remark

*Compared to its Type-I counterpart, Eq. (10), the Type-II variant of the iterative* 𝓁_2_ *method adaptively selects the parameter ϵ*_*n*_ *based on the previous iterations. The weight update rule is given by:*

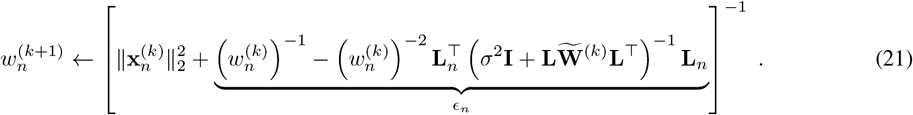

*Note that ϵ*_*n*_ *is an adaptively adjusted parameter that integrates information from both the noise covariance and prior weights. As highlighted in [21], this approach provides a principled mechanism to mitigate the risk of convergence to suboptimal local minima whenever possible*.

#### 2.3.3 Recovering Brain Sources with Free Orientation

The theory outlined so far assumes a fixed orientation model, where brain sources are constrained to a scalar voxel setting, typically perpendicular to the cortex surface. However, in real data, estimating surface normals is often challenging or undefined, particularly in volumetric reconstructions. To address this, we adopt a free orientation model, where each source is represented as an *O*-dimensional current vector (*O* = 3 for EEG, *O* = 2 for MEG), allowing for flexible dipole orientations. We set *O* to 2 for MEG as radial sources are not visible with MEG. It gives us only 2 tangential directions to possibly infer.

To accommodate free orientation sources, we utilize a full *O*-dimensional lead field matrix, denoted as: **L**^*O*^ ∈ ℝ^*M ×ON*^, where *N* is the number of voxels, and each **L**_*n*_ ∈ ℝ^*M ×O*^ represents the lead field matrix for the *n*-th voxel, capturing the *O* possible orientations of the source. Each column 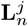 for *j* = 1, · · ·, *O* represents the observable scalp signal given a unit current dipole at voxel *n*, oriented along the *j*-th direction. Similarly the source activity at voxel *n* and time is modeled as 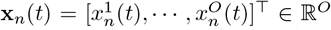, where 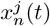 for *j* = 1, · · ·, *O* represents the current strength along each orientation at time *t*. For additional details on the update rules and mathematical derivations for the free orientation setting, please refer to the supplementary material.

**Table 1:**
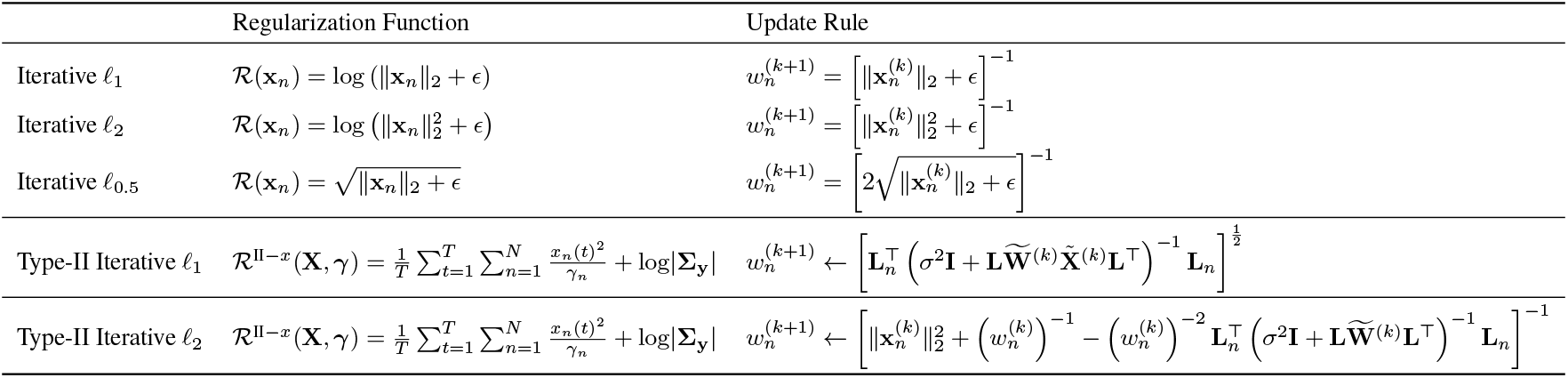
Update rules for iterative Type-I and Type-II methods described in Section 2. In each case, *ϵ* is a small constant added to ensure numerical stability. Also note that 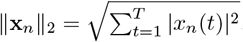.

### 2.4 Performance metrics

To rigorously assess the performance of BSI methods, four key performance metrics were employed: Earth Mover’s Distance (EMD), Euclidean distance, Mean Squared Error (MSE), and F1 score. All these metrics were computed between the estimated source localization and the corresponding ground truth. The mean values of these metrics across all subjects and runs are reported.

**EMD** [27, 6] is used to quantify the spatial localization accuracy. It is computed between the normalized values (to ensure that all elements are non-negative and sum to 1) of simulated and reconstructed sources, over the Euclidean distance (2-norm) space. It measures how much “work” is needed to move the reconstructed source points to match the true source points. Here, “work” is the distance each point needs to be moved times the amount of source strength being moved. When using Euclidean distance, EMD can be thought of as the minimum total distance required to move the reconstructed source to overlap with the true source. This metric has been considered in previous works on BSI such as [6] or [28].

**Euclidean distance** between each simulated source and the reconstructed source is calculated to evaluate the localization error. For a better intuitive interpretation of the Euclidean distance, we report it only for experiments with 1 non-zero source. The distance between the strongest estimated source with the true source is calculated.

**MSE** is calculated to evaluate the performance of the amplitude estimation, as a signed quantity. It is calculated as the squared differences between the reconstructed source amplitude values and the simulated source amplitude values, averaged over the entire time period and all sources.

**F1 score** is used to assess the recovery of the correct location of the true sources. It provides a balanced assessment of the ability of the BSI method to correctly identify brain sources while minimizing false positives and false negatives. A perfect recovery of true supports would yield an F1 score of 1.

**Residual noise** is calculated to measure the ability of a BSI method to effectively isolate and preserve genuine neural signals while minimizing the impact of noise. It is defined as

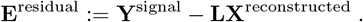

This quantity is used to calculate the **reconstructed noise level** reported in the figures, defined as the ratio of the Frobenius norm of **E**^residual^ and **Y**^signal^:

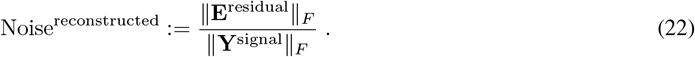

The true noise level, also reported in the figures, is defined as a ratio of **Y**^signal^ and **Y***/*^noise^(see 25):

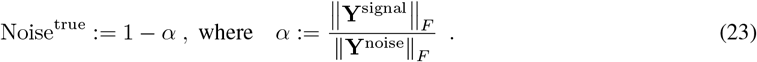

True noise level (simulated in the experiments) and reconstructed noise level serve as essential benchmark metrics. A true noise level value of 1 signifies that the signal is entirely composed of noise. Conversely, a true noise level value of 0 indicates a noiseless signal. The simulated noise level describes the amount of noise added to the signal during data generation, providing a consistent baseline for evaluations. Together with the simulated noise level, the reconstructed noise level gives insight into the amount of noise still present in the signal reconstructed by a BSI method as well as into the amount of genuine signal that might have been removed. Both quantities are scaled between 0 and 1, encompassing the entire range of possible values. This standardized scale ensures that different methods and approaches can be directly compared regardless of different ways in which BSI methods parameterize regularization strength/noise level, bridging a crucial gap in the field. Previously, the absence of such a unified metric hindered the comparability of various inverse solutions. Now, the performance of different methods can be assessed and benchmarked effectively.

### 2.5 Model Selection

#### 2.5.1 Grid Search

To uncover the optimal setting for each BSI method, we perform a comprehensive exploration of the regularization parameter in the BSI-Zoo. We systematically varied the regularization strength, *λ*, across a range of values, depending on the method - smooth linear inverse solution or sparse non-linear Bayesian learning solution. This exploration helps us understand how regularization affects the performance of source localization and identify configurations that work well.

#### 2.5.2 Spatial Cross-validation

In spatial cross-validation (CV), the data is split into training and test sets based on the MEG sensors [8, 29, 24]. We use “GridSearchCV” from the Scikit-learn software [30] with 5 CV folds. 80% of the sensors are assigned to the training set, denoted as 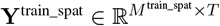, and the remaining 20% to the test set, denoted as 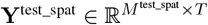.

The training sensors are used to fit the BSI model, employing the corresponding portion of the lead field matrix, **L**^train_spat^, exhaustively over *λ* values from the regularization parameter grid. The sources, represented by **X**^train_spat^ ∈ ℝ^*N ×T*^ and estimated from the fitted models, are subsequently mapped back to the sensor space. The MSE is then calculated on the heldout test sensors.

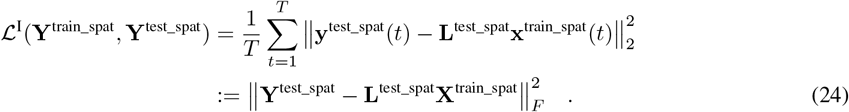

The value of *λ* that minimizes the MSE on the test sensors is chosen as the optimal regularization parameter.

**Remark**. *Note that spatial CV is not applicable to Type-II methods, as their loss function is based on Bregman divergence rather than Euclidean data fidelity, as observed in Eq. (24). Therefore, a metric other than MSE is needed to select the optimum value of λ that minimizes the Type-II loss. [24] introduced temporal CV for this purpose. Applying temporal CV to Type-II methods is a potential direction for future research, which we plan to pursue*.

## 3 Simulation Setup

In this section, through extensive simulations, we benchmark the performance of BSI methods at varying regularization strengths/estimated noise levels.

### 3.1 MEG Signal Simulation

#### 3.1.1 Forward Modeling

A forward model describes the linear relationship between the propagation of neuronal currents originating from brain source regions and the measurements recorded by E/MEG electrodes. Here we simulate MEG data with real lead field matrices. We employ realistic volume conductor models of human heads, which are taken from the publicly available CamCAN dataset [31]. We use individual T1 MRI scans of each subject to generate the corresponding lead field or forward model using the MNE software [32]. A minimum distance of 3 mm is set between the sources and the inner skull surface. We calculate the lead field matrices for each subject using the boundary element method (BEM), considering both fixed and free dipolar source orientations. These lead field matrices consist of 4686 sources and consider 204 MEG gradiometer channels.

#### 3.1.2 Source and Noise Model

We conducted simulations involving a limited number of active sources, which were randomly distributed across the cortex. The number of active sources remains fixed throughout the entire simulation. To represent the electrical neural activity of the brain sources, we sampled *T* = 10 data points from a standard normal distribution to model the evolving characteristics of each source over time. Next, we randomly distributed dipolar current sources across 4686 potential locations within the cortex, creating sparse source activation vectors denoted as **x**(*t*). These source activations, collectively denoted as **X** = [**x**(1), …, **x**(*T*)], were subsequently projected onto a set of 204 MEG sensors using the fixed-orientation lead field matrix **L** ∈ ℝ^204*×*4686^. This projection resulted in **Y**^signal^ = **LX**.

Next, we initialized either a diagonal or full-structural noise covariance matrix by drawing random samples from a Wishart distribution; with n degrees of freedom and a scale matrix equal to 10^−3^ times an identity matrix with n rows, where n equals the number of sensors. This ensures that the positive semi-definite (PSD) property is satisfied. The noise component, denoted by **Y**^noise^, was generated by sampling from a multivariate Gaussian distribution. This Gaussian distribution had a mean of zero and a covariance matrix that was derived from the aforementioned Wishart distribution. After the noise was normalized based on its Frobenius norm, we combined the signal and noise components to produce the pseudo-MEG signal as follows:

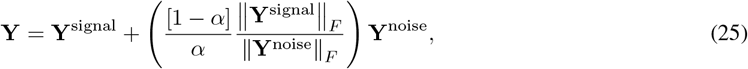

where, the parameter *α* is responsible for determining the signal-to-noise (SNR) ratio in the sensor space.

### 3.2 Parameter Initialisation

We simulate several experiments with varying benchmark parameters. For each simulation experiment, we test each BSI method on a grid of hyperparameters. The benchmark’s variable parameters and their values/ranges are specified in this section.

The number of non-zero sources (NNZ) was varied between 1, 2, 3 and 5. The number of time samples was set to *T* = 10. The parameter *α*, which determines the SNR in the simulated signal (see Eq.(25)) was varied from 0.99 to 0.01 (0.99, 0.9, 0.8, 0.7, 0.6, 0.5, 0.4, 0.3, 0.2, 0.1, 0.01). The regularisation parameter *λ* was varied across 10 values between 10^−3^ and 10^5^, scaled by magnitudes of 10 for eLORETA and ***γ***-MAP. For all other methods, 20 log-spaced *λ* values between 0.01 and 1 were tested. BSI methods were applied using lead field matrices accounting for both fixed and free dipole orientations. For the fixed orientation case, we perform 10 independent trials (*n*_fixed_ = 10) for each unique combination of subject, true SNR, regularization parameter, and NNZ. Each unique combination is referred to as an experiment run. Due to computational constraints, we limit our analysis to 1 independent trial (*n*_free_ = 1) in the free orientation case, for each experiment run. The random noise added varied across trials and source estimation and was performed for all experiment runs within each trial. In the experiments reported in this paper, we evaluate *K* = 4 distinct subjects to ensure methodological robustness. Hence, each unique simulation experiment is performed *n*_fixed/free_ time(s). Data generation was seeded for controlled and reproducible analyses. All parameter ranges were fixed after thorough exploration to balance between good coverage of the BSI method over reconstructed noise level and the practical constraints of computational resources.

After fitting BSI models, regularization parameters were converted into reconstructed noise levels using Eq.(22). For each performance metric, Gaussian process (GP) regression was used to extrapolate the attained performance across the entire plane spanned by true and reconstructed noise levels by aggregating information across subjects, trials, and different values of NNZ. The GP posterior distribution gives rise to an estimated mean performance surface as well as percentiles for each true/reconstructed noise level combination. In Figure 1 we report the 95th percentile, which can be interpreted as the estimated best performance that is attained or exceeded in 95% of cases. Mean GP performance is presented in supplementary Figure S1a. Results obtained from the free orientation experiments are reported in supplementary Section 3.

**Figure 1:**
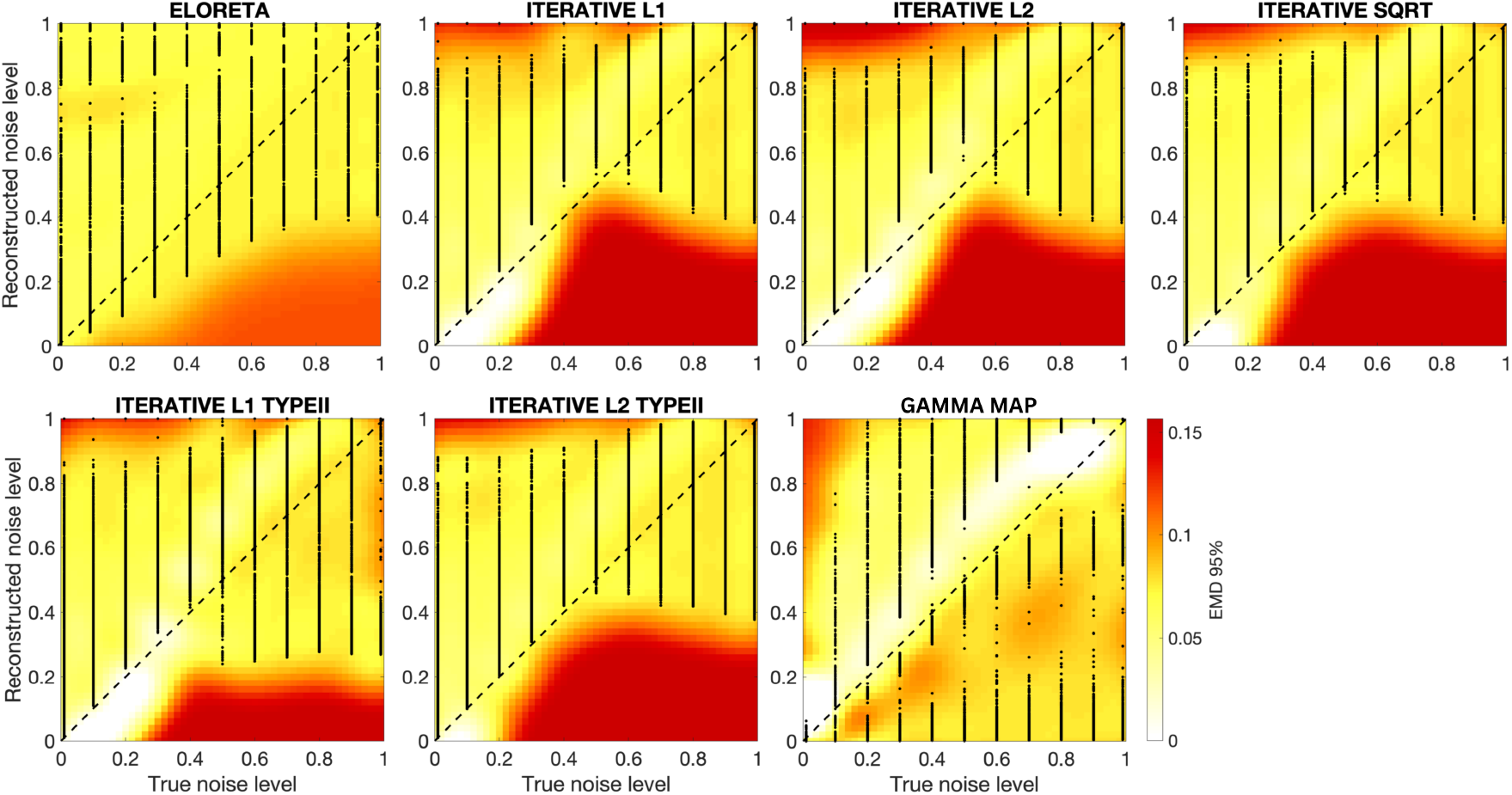
Source reconstruction performance of various Brain source imaging (BSI) methods as a function of true and reconstructed noise level. Performance is measured in terms of the Earth-Mover’s Distance (EMD), where lower EMD indicates better performance. Each heatmap illustrates the best performance that is attained or exceeded in 95% of cases (the 95th percentile of the posterior of a Gaussian process regression approximation of the EMD). Each black dot represents a simulation experiment. Across all BSI methods, it can be observed that better source reconstruction performance is generally observed when reconstructed noise levels match or slightly exceed true noise levels (are close or slightly above the diagonal), where better reconstruction is possible for lowest true and reconstructed SNR (lower left corner). Methods differ in terms of minimal attainable EMD but also with respect to the ranges of reconstructed noise levels within which performance is still competitive.

## 4 Simulation Results

We conducted an extensive evaluation of all BSI methods across varying levels of true and reconstructed noise levels, with knowledge of the ground truth source configurations to be reconstructed. The results discussed in this section are based on the fixed orientation simulations. Figure 1 showcases the diversity in performance, emphasizing the variability within and between methods resulting from different hyperparameters and simulation settings. Better performance (indicated by a lower EMD score) is attained when the reconstructed noise levels closely align with the true noise levels (along the diagonals of each subfigure). Generally, performance begins to drop with increasing true noise levels, and this deterioration varies uniquely for each method. Moreover, distinct performance outcomes emerge across the regularization grid, even at a consistent noise level.

Interestingly, our findings suggest that BSI methods tend to perform better with a moderate degree of underfitting (top left of the heatmap) than with overfitting (which typically results in a sharp performance drop), particularly at lower noise levels (below 50%). At higher noise levels, some degree of overfitting contributes to improved performance. This is further analyzed in Figure 4. However, this trend varies significantly depending on the specific BSI method, highlighting the method-dependent nature of the misspecification range within which a BSI method maintains optimal performance.

To align parameter choices with each BSI method’s intrinsic characteristics, we optimized the regularization parameter, *α*, using Spatial CV. In Figure 2, we compare the EMD performance achieved by spatial CV with the optimal performance attained by the respective method. Optimal performance represents the method’s best performance interpolated across all trials and experiment runs. Notably, spatial CV consistently yields performance closely approaching the optimal attainable performance.

**Figure 2:**
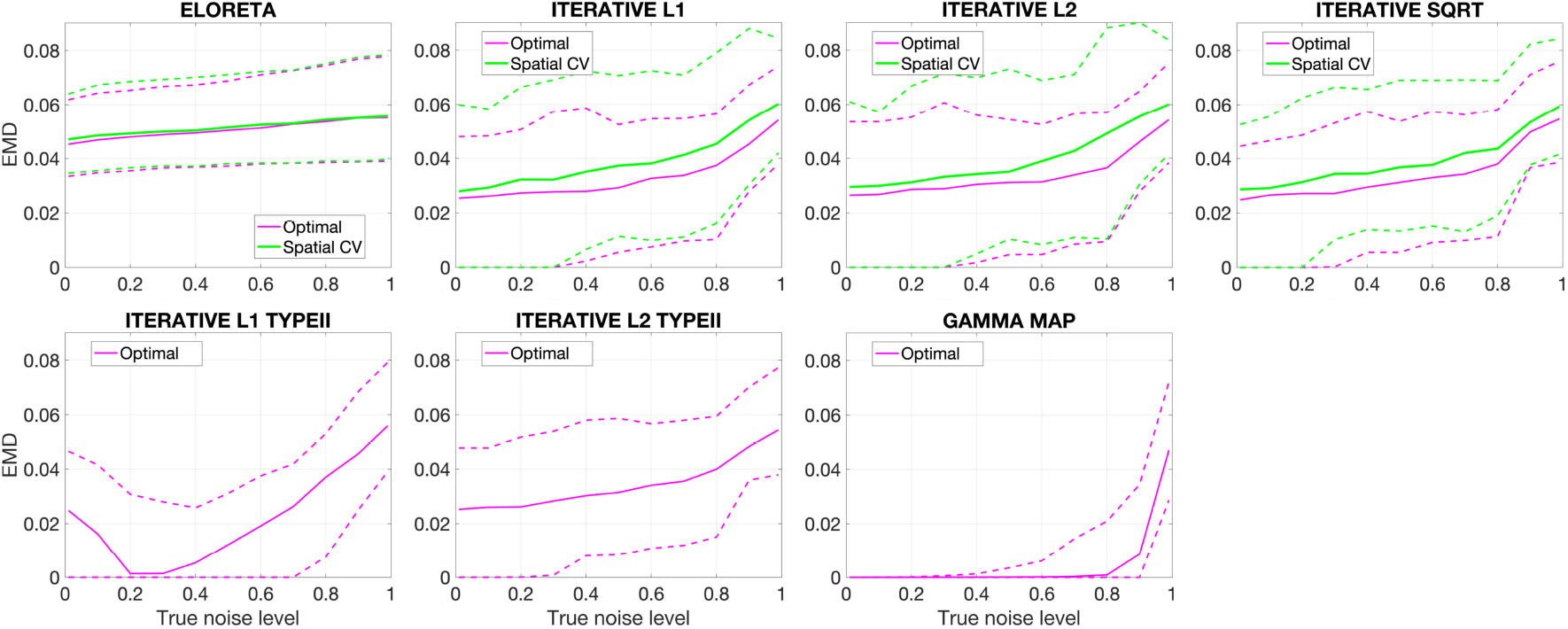
Comparison of optimal and spatial CV performance. Each plot illustrates a different BSI method’s optimal and spatial CV performance. Optimal performance represents the best result achieved across all possible choices of the regularization parameter of reconstructed noise level, while spatial CV performance is the result achieved from spatial cross-validation model selection. Solid lines denote the mean across experiments, while dotted lines denote the 95% confidence interval. Spatial CV consistently yields performance close to the optimal attainable performance across all noise levels.

Additionally, Figure 2 serves as a means to compare methods with each other in terms of their best achievable performance at a given noise level. Type-II methods, overall, demonstrate significantly superior performance, particularly at lower noise levels. ***γ***-MAP stands out by achieving the best optimal performance (lowest EMD) across all noise levels, showcasing its robustness. eLORETA exhibits consistent performance even as noise worsens. For all other methods, optimal performance declines with increasing noise levels.

Further analysis was conducted to assess the robustness of the methods. In Figure 3a, we present the global performance by averaging over all regularization and noise levels. This computation was carried out across all trials and experiment runs, as well as over the complete plane spanned by simulated and reconstructed noise level using the Gaussian process based inter- and extrapolation, integrating over the posterior means and 95th percentiles. This reiterates that Type-II methods demonstrate superior performance overall. Figure 3b illustrates the methods’ tolerance to noise level misspecification. We calculate the area (spanned by all combinations of true and reconstructed SNR) for which lower EMD values (better performance) were attained. This is calculated for three thresholds at the 25th, 50th, and 75th percentiles of attained EMD values (across all methods and experiments). Type-I methods exhibit greater susceptibility to misspecification, resulting in lower overall performance. eLORETA displays the highest tolerance to misspecification, as it shows the largest SNR area (integral over combinations of true and reconstructed noise levels) within which values below the median EMD across all experiments are attained. While fairly tolerant to misspecification, ***γ***-MAP exhibits the largest SNR area within which values below the 25th EMD percentile across all experiments are attained. Heatmaps (similar to Figure 1) thresholded by percentiles are presented in Supplementary Figure S1b. Overall, Type-II methods demonstrate higher peak performance but within a more narrow noise level misspecification range. In contrast, eLORETA, while not achieving the best performance, exhibits decent performance across a considerably broader noise misspecification range compared to the Type-II methods.

**Figure 3:**
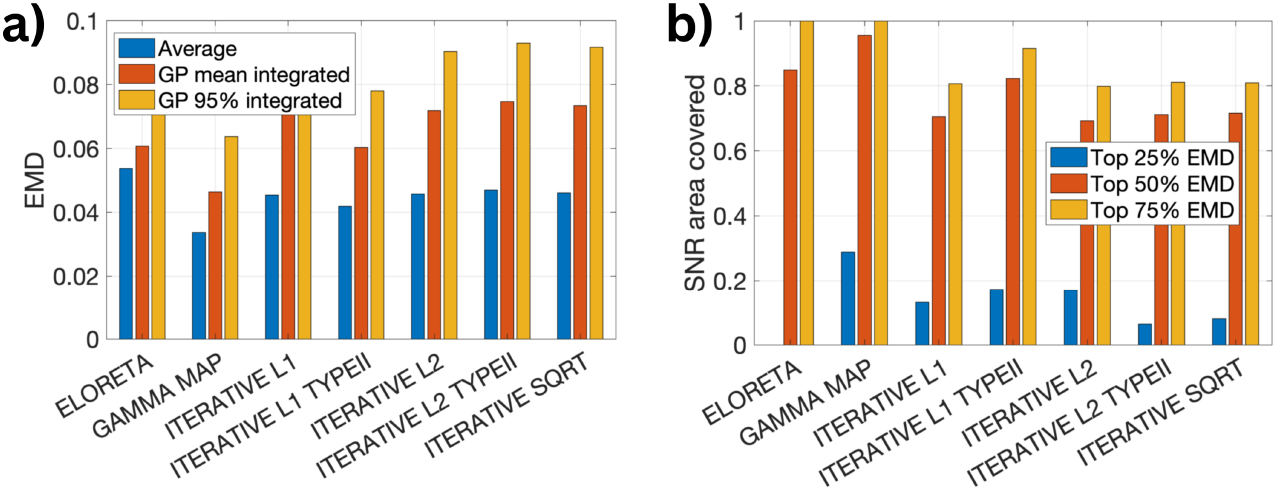
Global source reconstruction performance of BSI Methods. a) Mean EMD across all true and reconstructed noise levels, across simulated trials and experiments. Blue bars indicate the arithmetic mean over simulated trials and experiments. Red bars showcase the posterior mean of a Gaussian Process approximation. Yellow bars indicate the 95th percentile of the same Gaussian Process approximation. The GP integration yields higher averages than the arithmetic mean, which can be attributed to the fact that extrapolated performances at the boundary of the observed reconstructed SNR ranges enter the average. ***γ***-MAP demonstrates the best overall performance. b) Tolerance to noise misspecification. Three thresholds at the 25th, 50th, and 75th percentiles of attained EMD values across all methods and experiments were defined. For each method and threshold, the area (spanned by all combinations of true and reconstructed SNR) for which lower EMD values (better performance) were attained was integrated. Larger SNR areas covered suggest greater tolerance to noise misspecification. eLORETA is observed to be particularly lenient to misspecification as indicated by its high 50% performance bar, which indicates that this method attains EMD values better than the median almost throughout the entire SNR plane. ***γ***-MAP, though, attains EMD values below the 25th percentile for a larger number of combinations of true and reconstructed SNR than all other methods. Analysis in panel b) is based on the 95th percentile of a GP interpolation.

In Figure 4, we investigate the degree of underfitting required as the noise level varies. We assess the bias in the optimal reconstructed noise level, which represents the distance between the optimal reconstructed noise level and the true noise level, averaged across all trials and runs. Our findings reveal that a slight degree of underfitting is necessary to achieve optimal performance. Only at higher noise levels does overfitting lead to improved performance. Interestingly, this trend holds across all evaluated BSI methods in the fixed orientation case.

**Figure 4:**
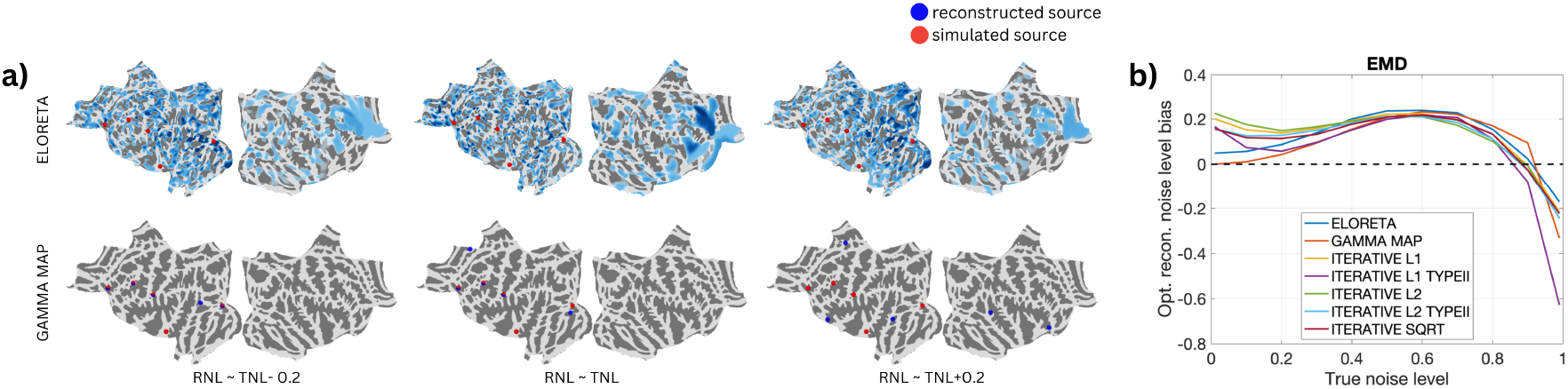
Bias and variance required with changing SNR. This figure illustrates the degree of underfitting or overfitting needed to attain optimal performance as the noise level increases. Panel a) illustrates the impact of different noise levels on source reconstruction by fixing the true noise level (TNL) at an intermediate value of 0.3. Source distributions across five degrees of reconstructed noise levels (RNL), TNL ± 0.2, TNL ± 0.1, TNL, are plotted to visually depict under- and overfitting effects. b) To quantify the degree of under/overfitting, we calculate the distance between the optimal and reconstructed noise levels across experimental trials. Positive values indicate underfitting, while negative values denote overfitting. The results from the fixed orientation simulations show that higher SNR levels require more underfitting, whereas overfitting becomes beneficial only at lower SNR levels.

Results from the experiments with free orientation of source exhibited similar trends across all analyses with reduced performance. Similar to the fixed orientation case, slight degrees of underfitting help achieve optimal performance. This can be seen in Supplementary Figure S6. Please refer to Supplementary Section 3 for other analysis figures on the free orientation experiments. Additional analysis for fixed orientation using other performance metrics is also reported in Supplementary Section 1.

## 5 Discussion

The accuracy of localizing neural sources is highly dependent upon modeling parameters. With several types of Brain Source Imaging (BSI) methods available, it is hard to formulate a one-fits-all rule to tune and evaluate parameter choices and further compare them. However, source localization is frequently compared solely on the best achieved performance, using the same dataset and under similar conditions (parameter settings).

In our study, we provide a comprehensive analysis of the performance of BSI methods across a wide range of conditions, varying noise levels, and regularization strengths, on semi-realistic MEG data. We investigated the resilience of EEG/MEG inverse solutions to noise level misspecification. By conducting extensive simulations and employing cross-validation techniques, we shed light on the varying optimal performance points of different BSI methods and ranges of regularisation strength under which the method is still reliable. We employ cross-validation techniques to identify these optimal performance points for each BSI method. Furthermore, we wrapped our approach in BSI-Zoo, an open-source Python package to facilitate standardized evaluations and comparisons of BSI methods.

Our findings yield valuable insights into the performance characteristics of BSI methods. Methods exhibit superior performance under high SNR levels. SNR level at which their performance begins to deteriorate varies for every method. Different BSI methods have different optimal performing points. This nuanced understanding derived from the benchmarking process contributes valuable insights for practitioners aiming to implement these methods in real-world scenarios, highlighting the importance of method-specific parameter calibration for achieving optimal outcomes.

Overall, ***γ***-MAP achieves the best performance across all noise levels. However, eLORETA is more forgiving to noise-level misspecification and exhibits consistent performance even as noise worsens. ***γ***-MAP is more susceptible to misspecification compared to eLORETA. Notably, underfitting tends to be preferable across all methods. Further, we demonstrate using cross-validation techniques for parameter tuning to attain optimal performance.

Based on our results, we recommend using spatial CV universally for estimating the noise level, particularly in combination with Type-I BSI methods. Our findings suggest that spatial CV may even outperform noise estimation from a baseline period, as the optimal noise level is achieved with slight underfitting rather than the true noise level. While baseline-based noise estimation is commonly employed in source localization studies, such as ERP analysis, our results indicate that relying on the oracle noise variance from a baseline may not be ideal. Instead, spatial CV inherently aligns more closely with the optimal regularization parameter, leading to better localization accuracy. Additionally, when no informed selection of the noise parameter is possible, eLORETA remains a robust choice, as it provides stable results without requiring fine-tuned noise level estimation.

The BSI-zoo toolbox enhances transparency, reproducibility, and collaboration in the field by offering a standardized and accessible platform for comparing BSI methods. We advocate its use to ensure rigorous evaluation and comparison of all new BSI methods. We suggest using MNE-Python for the convenient use of a BSI method.

While the study provides valuable insights, several limitations must be acknowledged. Firstly, only sparse sources were simulated, which restricts the generalizability of the findings to other source configurations. In real-world scenarios, neural activations are often more complex and may involve distributed or overlapping sources, which were not fully explored in this work. Secondly, the results from the free orientation setting are based on a single trial. This choice was made to reduce computational complexity but may limit the applicability of the results to scenarios where the orientation of the sources is more variable. Finally, we did not include mixed-norm methods specifically designed for the frequency domain. Moreover, beamforming methods, which rely on spatial filtering techniques, were not considered as they are not applicable to the chosen simulation setting.

In the future, we aim to expand our benchmark to include a wider variety of methods, such as noise learning techniques [24, 33, 34, 35]. This will also involve exploring different data generation techniques, including sources with autocorrelation structure [36, 37], and other types of noise, such as pink noise, and brain noise.

## Author Contributions

AN: Conceptualization, Data curation, Formal analysis, Investigation, Methodology, Project administration, Software, Validation, Visualization, Writing - original draft, Writing - review & editing. SH: Conceptualization, Methodology, Visualization, Writing - review & editing. AG: Conceptualization, Methodology, Data curation, Software, Writing - review & editing. AH: Conceptualization, Data curation, Formal analysis, Investigation, Methodology, Software, Validation, Supervision, Writing - original draft, Writing - review & editing.

## Acknowledgments

The code repository of this paper extensively utilized the MNE-Python open-source software package [38] for M/EEG data analysis and source localization. We gratefully acknowledge the MNE developers and contributors for their efforts in maintaining and advancing this tool.

The work conducted by AG was at Inria, AG is presently employed by Meta Platforms. All the datasets used for this work were not accessed or processed on Meta’s compute infrastructure.

## Data Availability Statement

The benchmark experiment results for this study can be found in the BSI-Zoo repository: https://github.com/braindatalab/BSI-Zoo/tree/master/bsi_zoo/data.

## Footnotes

1 https://github.com/braindatalab/BSI-Zoo

