## Supplementary Material for "How forgiving are M/EEG inverse solutions to noise level misspecification? An excursion into the BSI-Zoo"

#### 1 Additional results from fixed orientation simulations

This section presents the results from fixed orientation simulations, utilizing additional metrics to evaluate model performance across different SNR levels comprehensively.

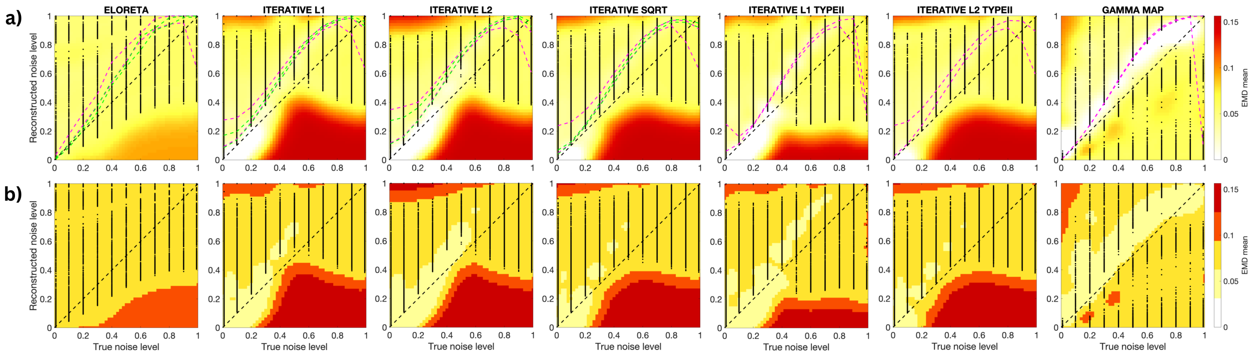


Figure S1: **Source reconstruction performance evaluated via Earth-Mover's Distance (EMD) across varying noise thresholds and confidence intervals for fixed orientation simulations.** Panel (a) shows the mean EMD with pink lines indicating the upper and lower bounds from the optimal performance and green lines marking the upper and lower bounds from cross-validation (CV). In panel (b), EMD is thresholded to visualize performance at four levels corresponding to 80%, 60%, 40%, and 20% of maximal loss (representing the least favorable performance). Lower EMD indicates better performance, with thresholded levels providing insight into the impact of noise on source reconstruction accuracy.

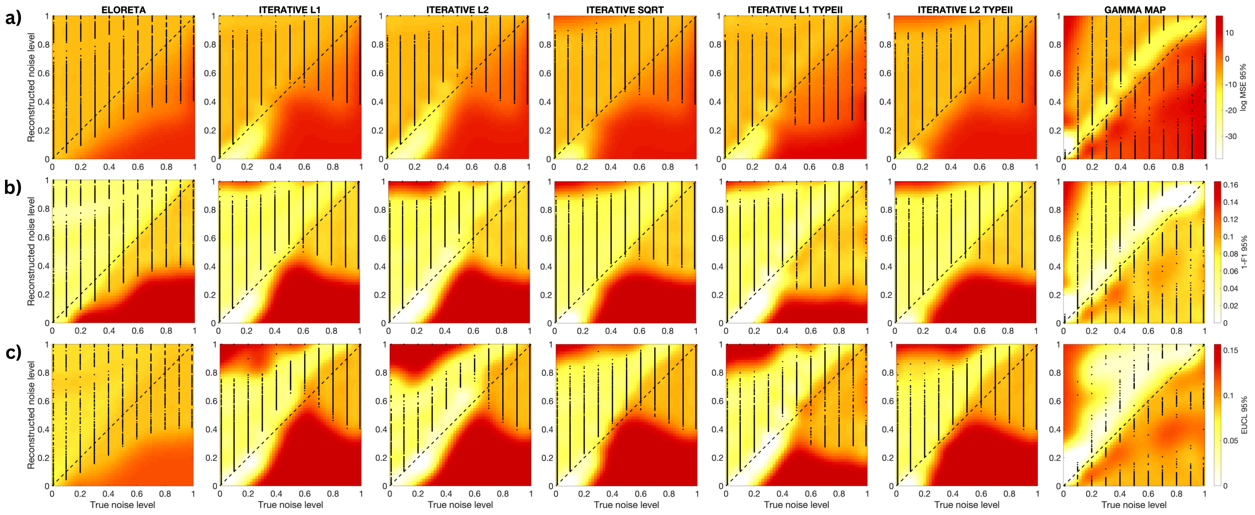


Figure S2: **Source reconstruction performance across metrics at the 95th percentile confidence level for fixed orientation simulations.** Panel (a) displays the log mean squared error (MSE), where lower values indicate better accuracy in source estimation. Panel (b) presents the F1 score, with higher scores reflecting improved precision-recall balance in source detection. Panel (c) shows the Euclidean distance (EUCL) for a single source, with lower distances indicating more precise localization. Each heatmap illustrates metric-specific performance trends at the 95% confidence threshold across varying noise and reconstruction levels.

#### 2 Recovering Brain Sources with Free Orientation

The theoretical derivations presented in the main paper assumed a “fixed-orientation” source model by considering a scalar voxel setting. In this scenario, the orientations of dipolar brain sources are presumed to be perpendicular to the cortex surface, necessitating estimation solely of the scalar deflection along the fixed orientation for each source. However, in real data, this assumption is often unrealistic, as surface normals may be difficult to estimate or even undefined, particularly in volumetric reconstructions. To address this, we adopt a “free-orientation” model, where each source is modeled as a full $O$-dimensional current vector, allowing it to flexibly align with the actual neural currents.

##### S2.1 Representing the Full 3D Lead Field

Initially, to accommodate the representation of free-orientation sources, the forward model is formulated in full 3D coordinates $(x, y, z)$, where each voxel $n$ is associated with a 3D dipole moment. The full 3D lead filed matrix is defined as: $\mathbf{L}^{3\mathrm{D}} \in \mathbb{R}^{M \times 3N}$ , structured as: $\mathbf{L}^{3\mathrm{D}} = [\mathbf{L}_1, \ldots, \mathbf{L}_N]$ , where $N$ represents the number of voxels, and each sub-matrix $\mathbf{L}_n = [\mathbf{L}_n^x, \mathbf{L}_n^y, \mathbf{L}_n^z] \in \mathbb{R}^{M \times 3}$ describes the lead field for voxel $n$ along its three Cartesian orientations. Each column $\mathbf{L}_n^j$ of $\mathbf{L}_n$ represents the measured signal at the scalp when a unit current dipole is positioned at voxel $n$ with an orientation along $j$, where $j \in x, y, z$.

The corresponding source activity is modeled as $\mathbf{x}_n(t) = [x_n^x(t), x_n^y(t), x_n^z(t)]^\top \in \mathbb{R}^{3 \times 1}$, where $\mathbf{x}_n(t)$ represents the time-dependent activity of the sources across its three spatial orientations. The generative model for the sensor data at time $t$ is:

$$\mathbf{y}(t) = \mathbf{L}^{3\mathrm{D}}\mathbf{x}(t) + \mathbf{e}(t) = \sum_{n=1}^{N} \mathbf{L}_n \mathbf{x}_n(t) + \mathbf{e}(t) \ . \tag{S1}$$

##### S2.2 Dimensionality Reduction via SVD

While the full 3D lead field provides a complete representation, in practice, not all three orientations per voxel contribute equally to the measured signal. Many studies reduce the dimensionality by selecting only the dominant orientations per voxel using singular value decomposition (SVD).

Applying SVD to the lead field matrix, we retain the first $O$ components per voxel, leading to a reduced $O$-dimensional lead field: $\mathbf{L}^O \in \mathbb{R}^{M \times ON}$, where $O$ represents the number of retained orientations per voxel, set as:

- $O = 3$ for EEG, retaining all three Cartesian components.
- $O = 2$ for MEG, where only two dominant components per voxel contribute to the measured field.

Thus, after SVD processing, each voxel $n$ is associated with a lead field: $\mathbf{L}_n = [\mathbf{L}_n^1, \ldots, \mathbf{L}_n^O] \in \mathbb{R}^{M \times O}$, where $\mathbf{L}_n^j$ represents the retained lead field components for orientation $j$ $(j = 1, \ldots, O)$. We apply SVD to the original 3D source activity across all time points, where the entire time series matrix for each voxel $n$ is represented as $\mathbf{X}_n \in \mathbb{R}^{3 \times T}$. This matrix is decomposed using SVD as follows:

$$\mathbf{X}_n = \mathbf{U}_n \mathbf{S}_n \mathbf{V}_n^\top , \tag{S2}$$

where $\mathbf{U}_n \in \mathbb{R}^{3 \times 3}$ is an orthonormal matrix, $\mathbf{S}_n \in \mathbb{R}^{3 \times 3}$ is a diagonal matrix containing the singular values, which represent the standard deviations of the SVD components in $\mathbf{V}_n$, and $\mathbf{V}_n \in \mathbb{R}^{T \times 3}$ is an orthonormal matrix representing the temporal structure of the signal.

By selecting the first $O$ dominant components per voxel, the reduced representation of the source activity is given by:

$$\tilde{\mathbf{x}}_n(t) = \mathbf{S}_n(1:O, 1:O)\mathbf{V}_n(t, 1:O)^\top \ . \tag{S3}$$

Note that the above equation uses a pseudo-notation where indexing follows MATLAB-style conventions, representing the selection of the first $O$ components $\mathbf{S}_n$ and corresponding rows from $\mathbf{V}_n$. This operation ensures that the retained components preserve the variance while reducing the dimensionality.

With this modification, the generative model is updated to:

$$\mathbf{y}(t) = \mathbf{L}^O \tilde{\mathbf{x}}(t) + \mathbf{e}(t) = \sum_{n=1}^{N} \mathbf{L}_n \tilde{\mathbf{x}}_n(t) + \mathbf{e}(t) \ . \tag{S4}$$

where $\tilde{\mathbf{x}}(t)$ is the reduced-dimensional source activity after SVD.

Note that for simplicity and consistency with the fixed orientation case, we continue to use the notation $\mathbf{x}_n(t)$, instead of $\tilde{\mathbf{x}}_n(t)$, to refer to the source activity after SVD dimensionality reduction. However, it should be noted that in the free orientation framework, this refers to the reduced-dimensional representation obtained through the SVD process.

##### S2.3 eLORETA for Brain Sources with Free Orientation

For free orientation sources, each source is represented as an $O$-dimensional current vector, spanning multiple time samples. Specifically, $\mathbf{x}_n$ is an $O \times T$ matrix that models the activity of source $n$ across $O$ orientations over $T$ time points, where $O = 3$ for EEG and $O = 2$ for MEG. Unlike the fixed orientation case, where each source is constrained to a single direction (e.g., perpendicular to the cortical surface), the free orientation framework allows sources to assume flexible orientations. To handle this increased complexity, eLORETA introduces orientation-specific priors and an iterative regularization process that balances the contributions of all dipole components while compensating for depth bias.

1. **Dipole Orientation Representation:** Each source $n$ is represented by an $O$-dimensional vector $\mathbf{x}_n(t) \in \mathbb{R}^O$, corresponding to the retained orientations per voxel at time $t$. This requires a $O \times O$ regularization matrix $\mathbf{R}_n$ for each source, which incorporates orientation-specific priors.
2. **Regularization Matrix Update:** The iterative update of $\mathbf{R}_n$ accounts for the coupling between the dipole components and employs a matrix inverse square root operation instead of a scalar operation.
3. **Regularization Term:** The scalar norm in fixed orientation case is replace with an orientation-aware regularization term, given by: $\lambda \sum_{n=1}^{N} \mathbf{x}_n(t)^\top \mathbf{R}_n \mathbf{x}_n(t)$, ensuring depth and orientation biases are jointly addressed.

In the free orientation setting, the eLORETA objective function is formulated as:

$$\mathbf{X}^{\text{eLORETA}} := \arg\min_{\mathbf{X}} \frac{1}{T} \sum_{t=1}^{T} \|\mathbf{y}(t) - \mathbf{L}\mathbf{x}(t)\|_2^2 + \lambda \sum_{t=1}^{T} \sum_{n=1}^{N} \mathbf{x}_n(t)^\top \mathbf{R}_n \mathbf{x}_n(t), \tag{S5}$$

$$= \arg\min_{\mathbf{X}} \|\mathbf{Y} - \mathbf{L}\mathbf{X}\|_F^2 + \lambda \,\mathrm{tr}(\mathbf{X}^\top \mathbf{R}\mathbf{X}) \tag{S6}$$

where $\lambda > 0$ is the regularization parameter controlling the balance between fitting the observed data and enforcing the prior, and $\mathbf{x}_n(t) \in \mathbb{R}^O$ represents the $O$-dimensional source activity at voxel $n$ at time $t$, where $O = 3$ for EEG and $O = 2$ for MEG. The term $\sum_{t=1}^{T} \sum_{n=1}^{N} \mathbf{x}_n(t)^\top \mathbf{R}_n \mathbf{x}_n(t)$ penalizes the energy of the dipole components, ensuring consistent contributions from all sources and mitigating depth bias.

Similar to fixed orientation setting, eLORETA follows an iterative approach to estimate the regularization matrices $\mathbf{R}_n$, which adaptively balance the contributions of different orientations while compensating for depth bias. As in the fixed orientation case, the lead field matrix $\mathbf{L}^O$ is pre-whitened to account for sensor noise, resulting in $\mathbf{G} = \mathbf{W}\mathbf{L}^O$, where $\mathbf{W}$ is the whitening matrix derived from the noise covariance matrix. Next, for each source $n$, the orientation-specific regularization matrix $\mathbf{R}_n$ is initialized as an identity matrix: $\mathbf{R}_n^{(0)} = \mathbf{I}_O,\ \forall n = 1, \dots, N$. This ensures that all sources initially contribute equally across their respective orientations, allowing the iterative process to refine the depth and orientation compensation. Then, at each iteration $k$, the orientation-specific regularization matrix $\mathbf{R}_n \in \mathbb{R}^{O \times O}$ is updated using an adaptive reweighting scheme:

$$\mathbf{R}_n^{(k+1)} = \left( \sum_{t=1}^{T} \mathbf{G}_n^\top \mathbf{H}^\top \mathbf{H} \mathbf{G}_n + \epsilon \mathbf{I} \right)^{-1/2}, \tag{S7}$$

where $\mathbf{G}_n$ is the contribution of the lead field matrix for source $n$, $\mathbf{H}$ is derived from the singular value decomposition (SVD) of $\mathbf{G}\mathbf{R}\mathbf{G}^\top$, and $\epsilon$ ensures numerical stability.

Convergence is determined by monitoring the relative change in $\mathbf{R}$ between iterations:

$$\delta := \frac{\|\mathbf{R}^{(k+1)} - \mathbf{R}^{(k)}\|_F}{\|\mathbf{R}^{(k)}\|_F}. \tag{S8}$$

The iterations stop when $\delta < \text{threshold}$, where the threshold is a small predefined value.

After convergence, the eLORETA kernel is computed as:

$$\mathbf{K} = \mathbf{R}^{1/2} \mathbf{G}^\top (\mathbf{G}\mathbf{R}\mathbf{G}^\top + \lambda \mathbf{I})^{-1}. \tag{S9}$$

Finally, the source estimates are obtained by applying the kernel to the pre-whitened sensor measurements:

$$\mathbf{X} = \mathbf{K}\mathbf{Y}. \tag{S10}$$

##### S2.4 Sparse Type-I Iterative Methods for Brain Sources with Free Orientation

As introduced earlier, for free orientation sources, each voxel is modeled as an $O$-dimensional current vector, where $O$ represents the number of orientations per voxel ($O = 3$ for EEG and $O = 2$ for MEG). Unlike the fixed orientation case, where a single weight per source is sufficient, in the free orientation setting, a key challenge is ensuring that all $O$ orientations within a voxel share a common sparsity pattern. This prevents different orientations from being treated independently, which would lead to inconsistent activation patterns across dipole components.

To enforce a consistent sparsity structure across orientations, a single regularization weight $w_n^{(k+1)}$ is assigned per voxel, rather than per orientation. This ensures that all dipole components within a voxel experience the same level of shrinkage, maintaining coherence in the estimated source activity. The regularization functions $\mathcal{R}(\mathbf{x}_n)$ and iterative updates $w_n^{(k+1)}$ for sparse Type-I iterative methods in the free orientation setting are formulated as follows:

- **Iterative** $\ell_1$ [1]:

$$\mathcal{R}(\mathbf{x}_n) = \log\left(\|\mathbf{x}_n\|_F + \epsilon\right), \quad w_n^{(k+1)} = \left[\|\mathbf{x}_n^{(k)}\|_F + \epsilon\right]^{-1}, \tag{S11}$$

  where $\|\mathbf{x}_n\|_F = \sqrt{\sum_{j=1}^{O}\sum_{t=1}^{T}|x_{n,j}(t)|^2}$ represents the Frobenius norm of the $O$-dimensional source activity over time and orientations, ensuring that all dipole components in a voxel share the same sparsity pattern.

- **Iterative** $\ell_2$ [1]:

$$\mathcal{R}(\mathbf{x}_n) = \log\left(\|\mathbf{x}_n\|_F^2 + \epsilon\right), \quad w_n^{(k+1)} = \left[\|\mathbf{x}_n^{(k)}\|_F^2 + \epsilon\right]^{-1}, \tag{S12}$$

  where $\|\mathbf{x}_n\|_F^2 = \sum_{j=1}^{O}\sum_{t=1}^{T}|x_{n,j}(t)|^2$.

- **Iterative** $\ell_{0.5}$ [2]:

$$\mathcal{R}(\mathbf{x}_n) = \sqrt{\|\mathbf{x}_n\|_F + \epsilon}, \quad w_n^{(k+1)} = \left[2\sqrt{\|\mathbf{x}_n^{(k)}\|_F + \epsilon}\right]^{-1}, \tag{S13}$$

  which is equivalent to

$$\mathcal{R}(\mathbf{x}_n) = \sqrt{\sqrt{\sum_{j=1}^{O}\sum_{t=1}^{T}|x_{n,j}(t)|^2} + \epsilon}, \quad w_n^{(k+1)} = \left[2\sqrt{\sqrt{\sum_{j=1}^{O}\sum_{t=1}^{T}\left|x_{n,j}^{(k)}(t)\right|^2} + \epsilon}\right]^{-1}.$$

In each free orientation formulation, $\|\mathbf{x}_n\|_F$ represents the Frobenius norm of the $O$-dimensional source activity over both time and orientation, ensuring that all the dipole components within a voxel contribute consistently to the regularization. The parameter$\epsilon$ ensures numerical stability in the regularization and weight update functions.

##### S2.5 Type-II Methods for Brain Sources with Free Orientation

In the Champagne or $\gamma$-MAP approach, free orientation sources are modeled by assigning a single variance parameter per source, shared across all its $O$ orientations. This ensures all orientations of a given source follow the same sparsity pattern, preventing inconsistencies that could arise if each orientation had an independent weight. The source covariance matrix in this case is defined as $\mathbf{\Gamma}^O = \mathrm{diag}(\boldsymbol{\gamma}^O) = [\gamma_1, \gamma_1, \ldots, \gamma_N, \gamma_N]^\top$, where each source $n$ has a single variance parameter $\gamma_n$, shared across its $O$ orientations.

Given this assumption, the prior distribution of the sources is given by: $\mathbf{x}(t) \sim \mathcal{N}(\mathbf{0}, \mathbf{\Gamma}^O)$, where $\mathbf{\Gamma}^O$ is defined as $ON \times ON$ block diagonal matrix expressed as

$$\mathbf{\Gamma}^O = \begin{bmatrix} \gamma_1\mathbf{I}_O & 0 & \cdots & 0 \\ 0 & \gamma_2\mathbf{I}_O & \cdots & 0 \\ \vdots & \vdots & \ddots & \vdots \\ 0 & 0 & \cdots & \gamma_N\mathbf{I}_O \end{bmatrix}, \tag{S14}$$

in which $\gamma_n\mathbf{I}_O$ is a prior variance $O \times O$ matrix of $\mathbf{x}_n$ and $\mathbf{I}_O$ is a $O \times O$ identity matrix.

The prior distribution $p(\mathbf{X}|\mathbf{\Gamma}^O)$ is then defined as

$$p(\mathbf{X}|\mathbf{\Gamma}^O) = \prod_{t=1}^{T}\mathcal{N}(\mathbf{x}(t)|\mathbf{0}, \mathbf{\Gamma}^O)) = \prod_{t=1}^{T}\prod_{n=1}^{N}\mathcal{N}(\mathbf{x}_n(t)|\mathbf{0}, \gamma_n\mathbf{I}_O), \tag{S15}$$

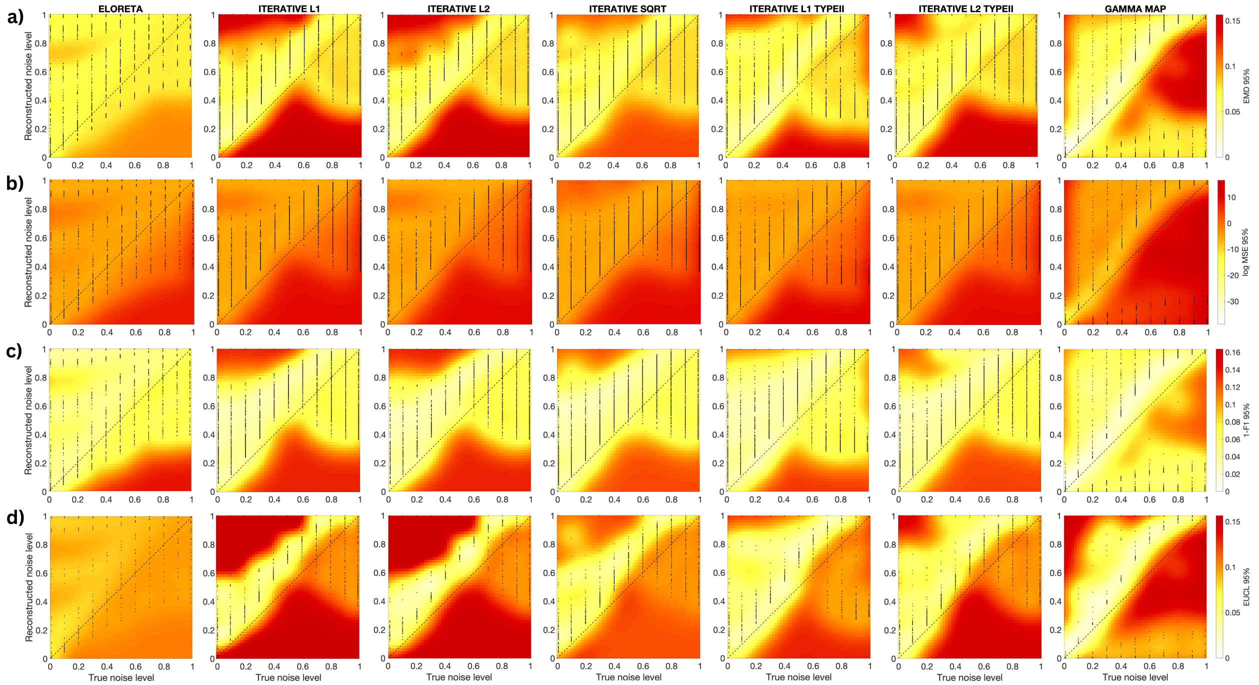


Figure S3: **Heatmaps of source reconstruction performance metrics at the 95th percentile confidence level for free orientation simulations.** Panel (a) shows the Earth Mover's Distance (EMD), (b) the mean squared error (MSE), (c) the F1 score, and (d) the Euclidean distance (EUCL) for a single source. Each heatmap captures variations in metric-specific performance at the 95% confidence threshold across varying noise and reconstruction levels. A drop in overall performance compared to fixed orientation experiments is observed under the free orientation setting. Figures are based on results from a single trial per experiment configuration.

### 3 Results from free orientation simulations

This section presents results from free orientation simulations. In the results reported here, we evaluate K = 4 distinct subjects. We perform 1 trial for each combination of subject, true SNR, regularization parameter, and NNZ, with parameters varied similarly to the fixed orientation setting.

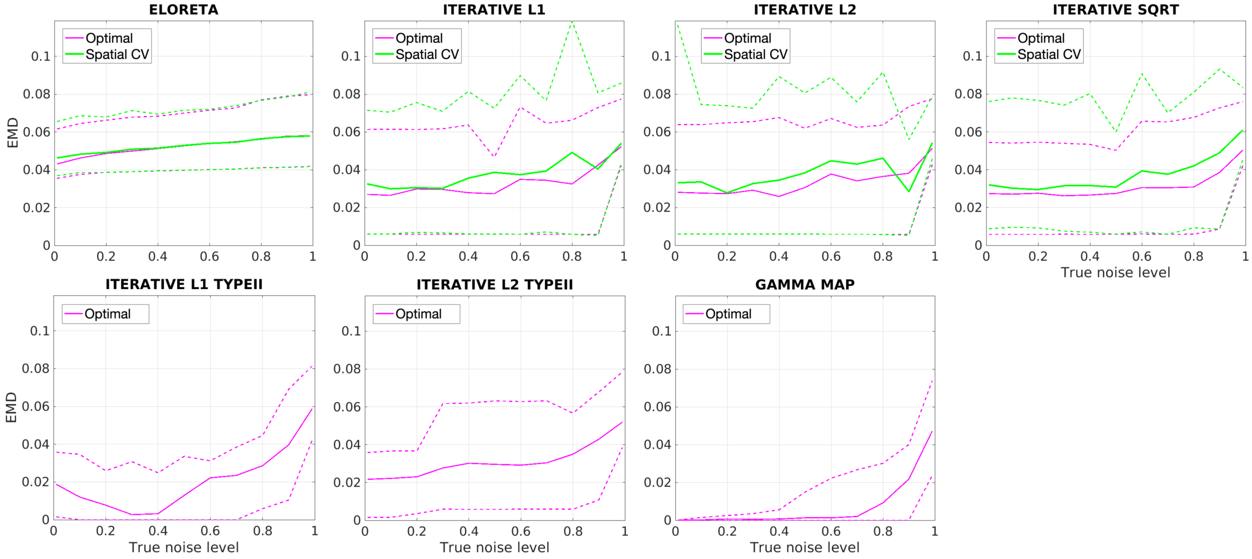


Figure S4: **Comparison of optimal and spatial CV performance in free orientation simulations.** Each plot displays the performance of optimal and spatial CV selections across noise levels, where optimal performance reflects the best outcome across all choices of the regularization parameter for reconstructed noise level, and spatial CV performance is derived from spatial cross-validation model selection. Solid lines indicate the mean across experiments, with dotted lines representing the 95% confidence interval. Spatial CV consistently achieves performance close to the optimal across varying noise levels even for free orientation case. Figures are based on results from a single trial per experiment configuration.

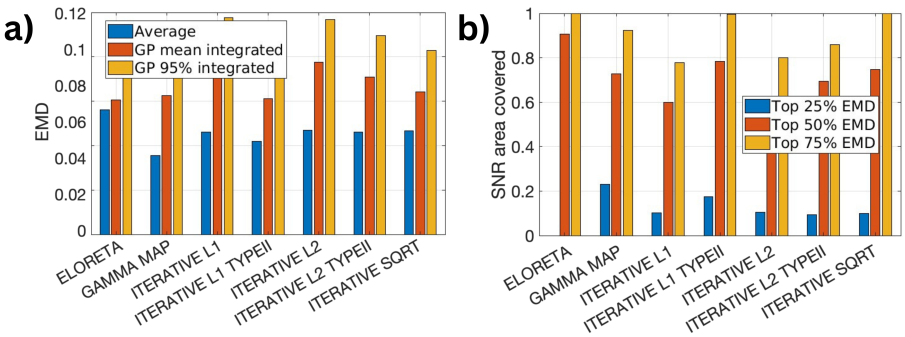


Figure S5: **Global source reconstruction performance in free orientation simulations.** (a) Mean Earth Mover's Distance (EMD) across all true and reconstructed noise levels, aggregated over trials and experiments. Blue bars represent the arithmetic mean, red bars indicate the posterior mean of a Gaussian Process approximation, and yellow bars show the 95th percentile of the same Gaussian Process. The Gaussian Process integration yields higher averages compared to the arithmetic mean due to extrapolated boundary performances in reconstructed SNR ranges. (b) Noise misspecification tolerance across methods, with thresholds set at the 25th, 50th, and 75th percentiles of EMD values. For each method and threshold, the area (spanning true and reconstructed SNR combinations) for which lower EMD values were achieved was integrated, with larger areas indicating greater tolerance to noise misspecification. This analysis provides insight into each method's robustness, with eLORETA showing strong tolerance as indicated by its extensive area under the 50% performance bar, while $\gamma$-MAP demonstrates high performance at stricter thresholds by attaining EMD values below the 25th percentile for a larger range of SNR combinations. Figures are based on results from a single trial per experiment configuration. (a) Mean Earth Mover's Distance (EMD) across all true and reconstructed noise levels, aggregated over trials and experiments. Blue bars represent the arithmetic mean, red bars indicate the posterior mean of a Gaussian Process approximation, and yellow bars show the 95th percentile of the same Gaussian Process. The Gaussian Process integration yields higher averages compared to the arithmetic mean due to extrapolated boundary performances in reconstructed SNR ranges. (b) Noise misspecification tolerance across methods, with thresholds set at the 25th, 50th, and 75th percentiles of EMD values. For each method and threshold, the area (spanning true and reconstructed SNR combinations) for which lower EMD values were achieved was integrated, with larger areas indicating greater tolerance to noise misspecification. This analysis provides insight into each method's robustness, with eLORETA showing strong tolerance as indicated by its extensive area under the 50% performance bar. $\gamma$-MAP demonstrates high performance at stricter thresholds by attaining EMD values below the 25th percentile for a larger range of SNR combinations. Figures are based on results from a single trial per experiment configuration.

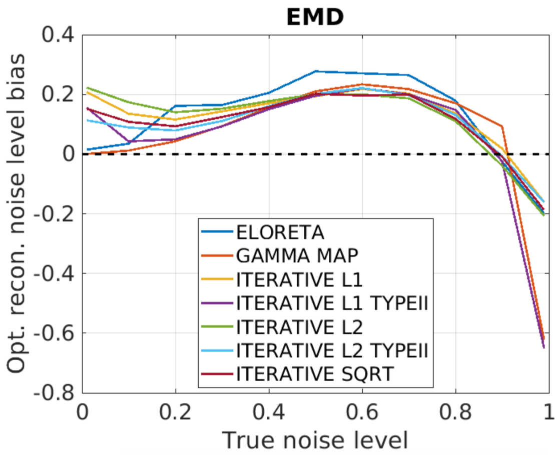


Figure S6: **Bias and variance across varying SNR in free orientation simulations.** To quantify the degree of underfitting or overfitting, we calculate the distance between the optimal and reconstructed noise levels across experimental trials. Positive values indicate underfitting, while negative values represent overfitting. Results show that underfitting is necessary to achieve optimal performance for all methods. The figure is based on results from a single trial per experiment configuration
